# A divergent CheW confers plasticity to nucleoid-associated chemosensory arrays

**DOI:** 10.1101/239202

**Authors:** Annick Guiseppi, Juan Jesus Vicente, Julien Herrou, Deborah Byrne, Aurelie Barneoud, Audrey Moine, Leon Espinosa, Marie-Jeanne Basse, Virginie Molle, Tâm Mignot, Philippe Roche, Emilia M.F. Mauriello

**Author notes:** These authors contributed equally (Annick Guiseppi, Juan Jesus Vicente). Correspondence to: Emilia M.F. Mauriello, Aix Marseille Univ, CNRS, LCB, 31 Chemin Joseph Aiguier, Marseille, 13402, France.

## Abstract

Chemosensory systems are highly organized signaling pathways that allow bacteria to adapt to environmental changes. The Frz chemosensory system from *M. xanthus* possesses two CheW-like proteins, FrzA (the core CheW) and FrzB. We found that FrzB does not interact with FrzE (the cognate CheA) as it lacks the amino acid region responsible for this interaction. FrzB, instead, acts upstream of FrzCD in the regulation of *M. xanthus* chemotaxis behaviors and activates the Frz pathway by allowing the formation and distribution of multiple chemosensory clusters on the nucleoid. These results, together, show that the lack of the CheA-interacting region in FrzB confers new functions to this small protein.

**AUTHOR SUMMARY:** Chemosensory systems are signaling complexes that are widespread in bacteria and allow the modulation of different cellular functions, such as taxis and development, in response to the environment. We show that the *Myxococcus xanthus* FrzB is a divergent CheW lacking the region involved in the interaction with the histidine kinase FrzE. Instead, it acts upstream of FrzCD to allow the formation of multiple distributed Frz chemosensory arrays at the nucleoid. The loss of the CheA-interacting region in FrzB might have been selected to confer plasticity to nucleoid-associated chemosensory systems. By unraveling a new accessory protein and its function, this work opens new insights into the knowledge of the regulatory potentials of bacterial chemosensory systems.

## INTRODUCTION

Chemosensory systems are specialized regulatory pathways that allow bacteria to perceive their external environment and respond with various cellular behaviors (1–3). In these systems, environmental signals are transduced inside the cells, initially by receptors called Methyl-accepting Chemotaxis Proteins (MCP). Most MCPs possess a transmembrane domain, but 14% are soluble proteins (4). Ligand-bound MCPs regulate the autophosphorylation of the histidine kinase CheA that, in turn, transfers phosphoryl groups to at least two response regulators: CheY which is the most downstream component of the pathway and CheB, which, together with CheR, constitutes the adaptation module. Che systems also include one or more versions of the docking protein CheW, which interacts with the C-terminal cytoplasmic tip of the MCP and with the P5 domain of CheA. CheW and CheA^P5^ are paralogs and are topologically similar to SH3 domains from eukaryotic scaffold proteins that also play a role in signal transduction. Such domains are characterized by two β-sheets (each composed of five β-strands) designated subdomain 1 and 2 (5–7). MCP: CheA^P5^: CheW interactions all involve subdomains 1 and 2. In particular, the β-strands 3 and 4 of subdomain 1 of CheA^P5^ interact with the β-strands 4 and 5 of subdomain 2 of CheW (8). The receptor cytoplasmic tip binds CheW or CheA at the hydrophobic junction between subdomain 1 and 2 (8).

High-resolution microscopy revealed that, intracellularly, Che proteins are organized in complex arrays running parallel to the bacterial inner membrane, with the MCP layer linked to a CheA/W baseplate (9,10). Transverse views of these Che arrays show ordered MCP hexagons networked by CheA-CheW rings. CheA-CheW rings contain alternating and interacting CheA^P5^ domains and CheWs. MCPs hexagons are constituted by six MCP trimers of dimers. One MCP in the dimer is connected with either a CheA^P5^ domain or a CheW, which allows connections between the MCP hexagons and the CheW-CheA^P5^ rings (11) (Fig. 1A). The presence of each of these three proteins is essential for the integrity of the system (12,13). This structure provides the typical signaling properties of chemotaxis systems, including signal amplification and sensitivity (14–17).

**Fig. 1.**
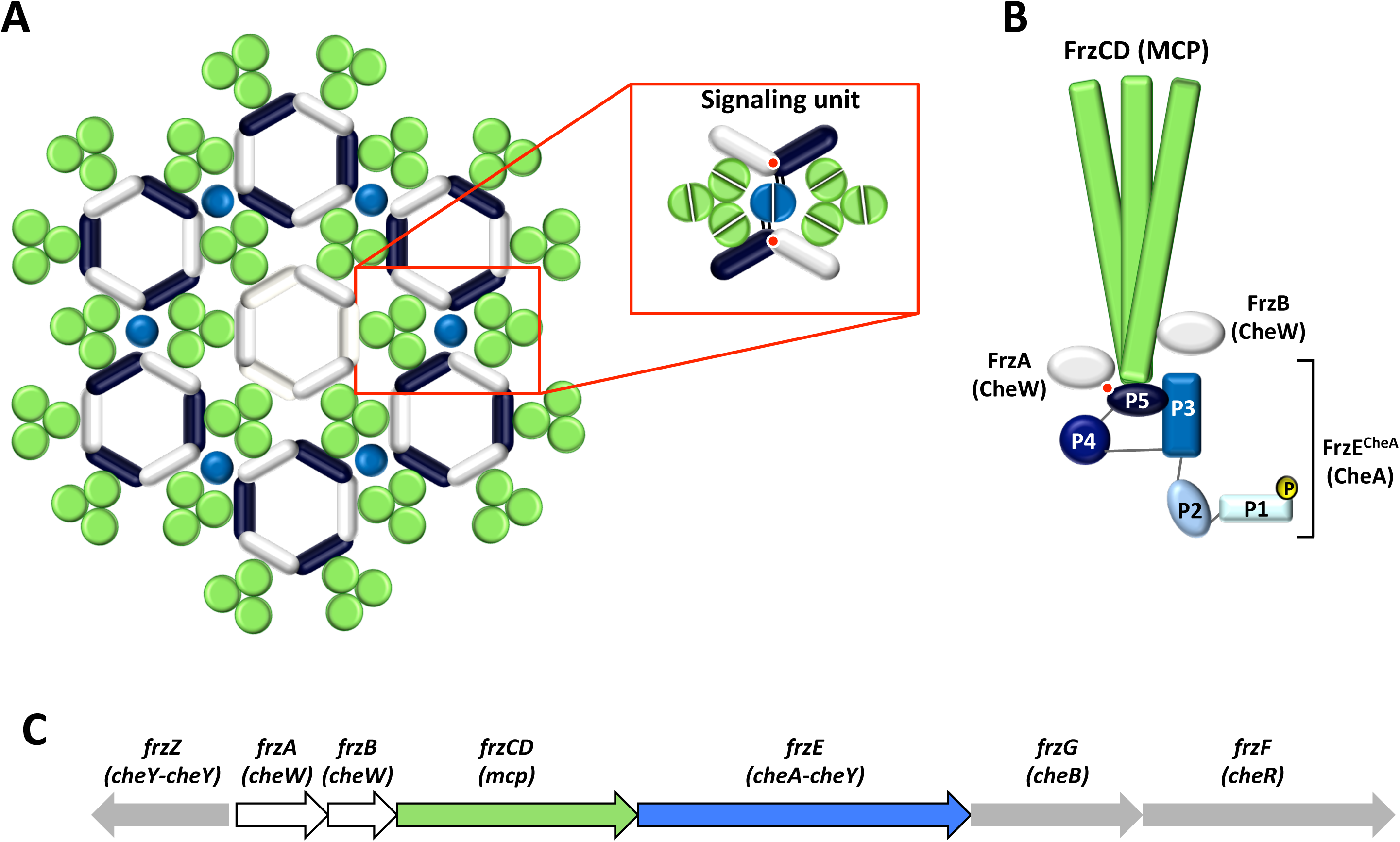
Schematic representation of the supramolecular organization of Che proteins. (A) MCP form trimers of dimers (each dimer is shown as a green circle), which, in turn, form hexagons connected with rings composed of the CheA-P5 domain (dark blue bars) and CheW (white bars). The light blue circles represent the CheA-P4 domain and, the red circles the interface between the β-strands 3 and 4 of subdomain 1 of CheA-P5 and the β-strands 4 and 5 of subdomain 2 of CheW. Rings containing six CheW proteins (shown at the center of the array) might serve to modulate the stability and activation of the system. A signaling unit is represented in the red box (Adapted from (43)). (B) FrzCD, FrzE^CheA^, FrzA and FrzB proteins organization depicted by homology with Che proteins. (C) Schematic representation of the *frz* operon.

The gliding bacterium *M. xanthus* uses the cytoplasmic Frz signal transduction pathway to bias the frequency at which cells change their direction of movement in response to environmental signals (18). This behavior allows cells to glide towards favorable directions or away from toxic compounds. The Frz system induces cell reversal by promoting the coordinated inversion of the cell polarity of the two *M. xanthus* motility systems: the Social (S) motility system powered by polar Type IV Pili and the Adventurous (A) motility powered by polarly assembled Agl-Glt complexes (19–21). The regulation of the motility systems by the Frz pathway is essential for cells to enter a developmental program that leads to the formation of multicellular fruiting bodies. The Frz core is composed of a cytoplasmic MCP (FrzCD), a CheA (FrzE) and a CheW (FrzA) (22) (Fig. 1B), encoded by a single operon (Fig. 1C). In the absence of any of these three proteins, cells display drastically reduced reversal frequencies and are no longer able to respond to isoamyl alcohol (IAA), a Frz activator (20,22). The Frz system also includes a second CheW-like protein, FrzB, described as an accessory because while in its absence cells show phenotypes similar to those caused by the deletion of core proteins, *ΔfrzB* cells are still able to respond to IAA with increased reversal frequencies (20,22) (Fig. 1B).

In this paper, we show that FrzB is a divergent CheW lacking the β-strands involved in the interaction with the histidine kinase FrzE (β4 and β5). Strikingly, the introduction of β4 and β5 from FrzA to FrzB converts the latter into a canonical CheW. Instead of mediating signal transduction from the FrzCD receptor to FrzE, FrzB acts upstream of FrzCD to allow the formation of multiple distributed Frz chemosensory clusters at the nucleoid. These clusters collapse in a unique and less active array in the absence of FrzB. Finally, the loss of the CheA-interacting region in FrzB might have been selected to confer plasticity to nucleoid-associated chemosensory systems.

## RESULTS

### FrzB is upstream of the FrzCD receptor in the Frz signaling pathway

The first two genes of the *frz* operon encode the CheW-like proteins, FrzA and FrzB. *ΔfrzA* cells show the same motility and fruiting body formation phenotypes as *ΔfrzCD* and *ΔfrzE*, suggesting that FrzA is the core CheW of the Frz pathway (22,23) (Fig. 2A). The aberrant behaviors at the colony level are due to the inability of *ΔfrzA* strains to modulate single-cell reversal frequencies even in the presence of isoamyl alcohol (IAA), a known Frz activator (Fig. 2B) (20,22).

**Fig. 2.**
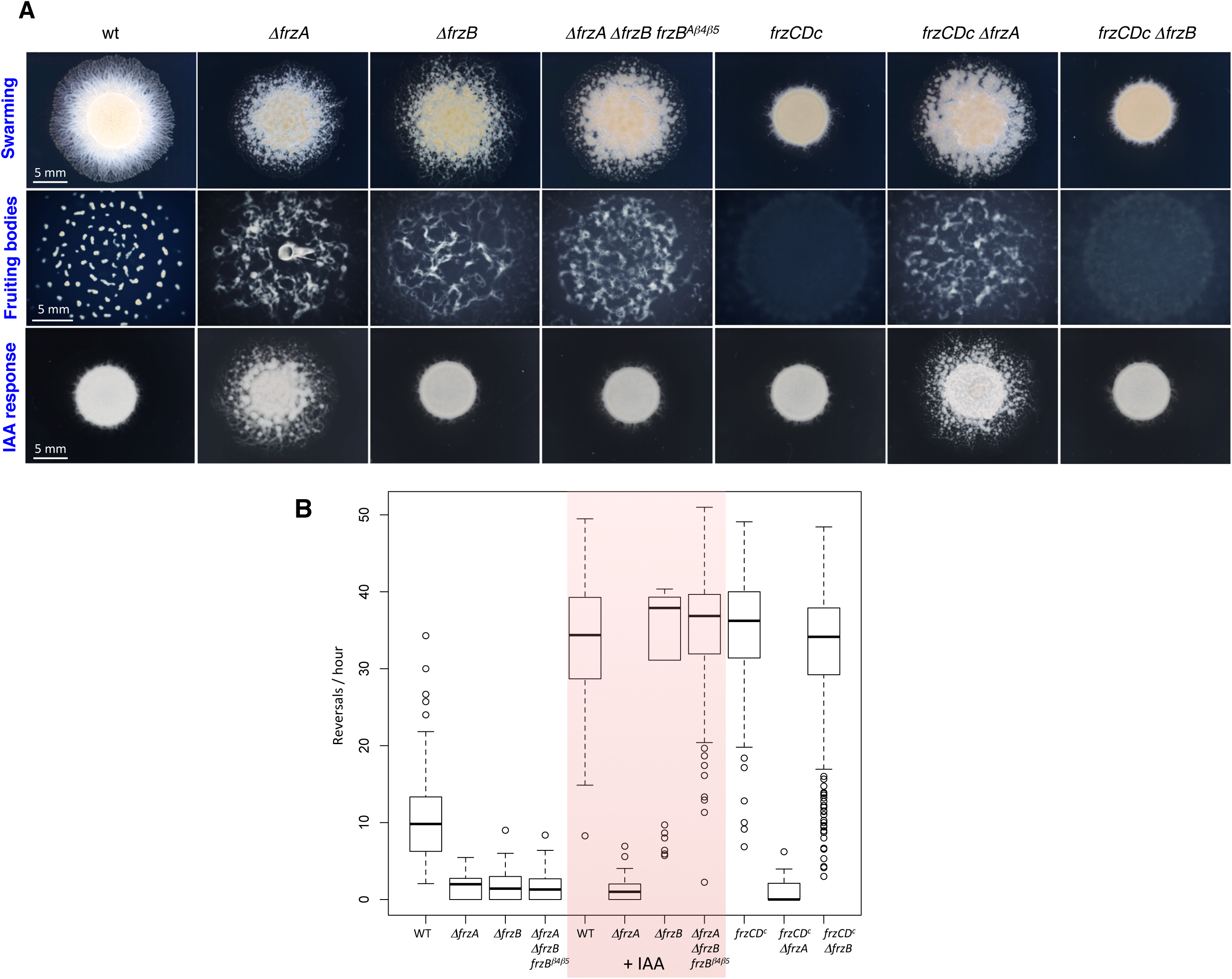
The effect of FrzB on the IAA response and the location of the *frzB* gene in the Frz pathway. (A) Motility and fruiting body formation phenotypes were photographed at 48h and 72h, respectively. (B) Box plots of reversal frequencies of single cells moving on agar pads supplemented or not with 0.15% IAA. The lower and upper boundaries of the boxes correspond to 25th and 75th percentiles, respectively. The median is shown as a line at the center of each box, and whiskers represents the 10th and 90th percentiles. For the reversal frequency measurements, approximately 500 cells from three to six biological replicates, were used. *frzCDc* corresponds to the allelic variant *frzCDΔ6-182*.

While a *frzB* deletion causes the same defects as *ΔfrzA* in the absence of IAA, IAA completely rescues the reversal frequency defect of *ΔfrzB* cells (Fig. 2B). This effect is detectable at both single-cell and colony level (Fig. 2) (22). These results show that signals from the FrzCD receptor can still be transduced to FrzE in the absence of FrzB and, thus, FrzB is not part of the Frz core. Moreover, a double *ΔfrzA ΔfrzB* phenocopies *ΔfrzA*, further suggesting the central role of FrzA and the accessory role of FrzB (Figure S1).

In order to determine the functional position of FrzB in the Frz pathway, we tested the epistasis of FrzB with respect to FrzCD. Because FrzCD is a central component of the Frz pathway, *ΔfrzB ΔfrzCD* and *ΔfrzA ΔfrzCD* double mutants showed the same aberrant motility and developmental phenotypes as *ΔfrzCD* (and *ΔfrzA*) at both single-cells and colony levels (Figure S1 and Fig. 2) (22). We thus took advantage of a *frzCD* allele, *frzCDc*, whose expression confers a constitutively hyper-reversing phenotype (Fig. 2) (Bustamante et al., 2004) and constructed *ΔfrzA frzCDc* and *ΔfrzB frzCDc* double mutants. The *ΔfrzA frzCDc* reversal frequencies and colony phenotypes were indistinguishable from those of *ΔfrzA* cells, confirming that FrzA is located downstream of the receptor in the signaling pathway (Fig. 2). On the other hand, *ΔfrzB frzCDc* phenotypes resembled those of *frzCDc* (Fig. 2), indicating that FrzB is upstream of FrzCD in the Frz signaling pathway and that its requirement is bypassed by a constitutive *frzCDc* mutation.

### FrzB does not interact with the downstream histidine kinase FrzE

Genetic data show that FrzB is upstream of FrzCD in the Frz regulatory pathway. We thus wanted to verify if FrzB could interact with FrzE. For this, we heterologously expressed and co-purified, using GST-affinity chromatography, GST-FrzA or GST-FrzB with either 6His-FrzCD or 6His-FrzE^CheA^. Both 6His-FrzCD and 6His-FrzE^CheA^ could be co-purified with GST-FrzA, suggesting that FrzA interacts with both FrzCD and FrzE^CheA^ (Fig. 3A). On the other hand, only 6His-FrzCD, but not 6His-FrzE^CheA^, could be co-purified with GST-FrzB, suggesting that FrzB only interacted with FrzCD but not with FrzE^CheA^ (Fig. 3A). We then wanted to test whether, by using high amounts of GST-FrzB, we could detect an interaction with immobilized 6His-FrzE^CheA^ in Biolayer interferometry (BLi) (24). While low amounts of GST-FrzB still produced an interaction with immobilized 6His-FrzCD (Kd = 0.7 μM), no interactions could be detected with immobilized 6His-FrzE1-658, even when we used high amounts of GST-FrzB (Fig. 3B).

**Fig. 3.**
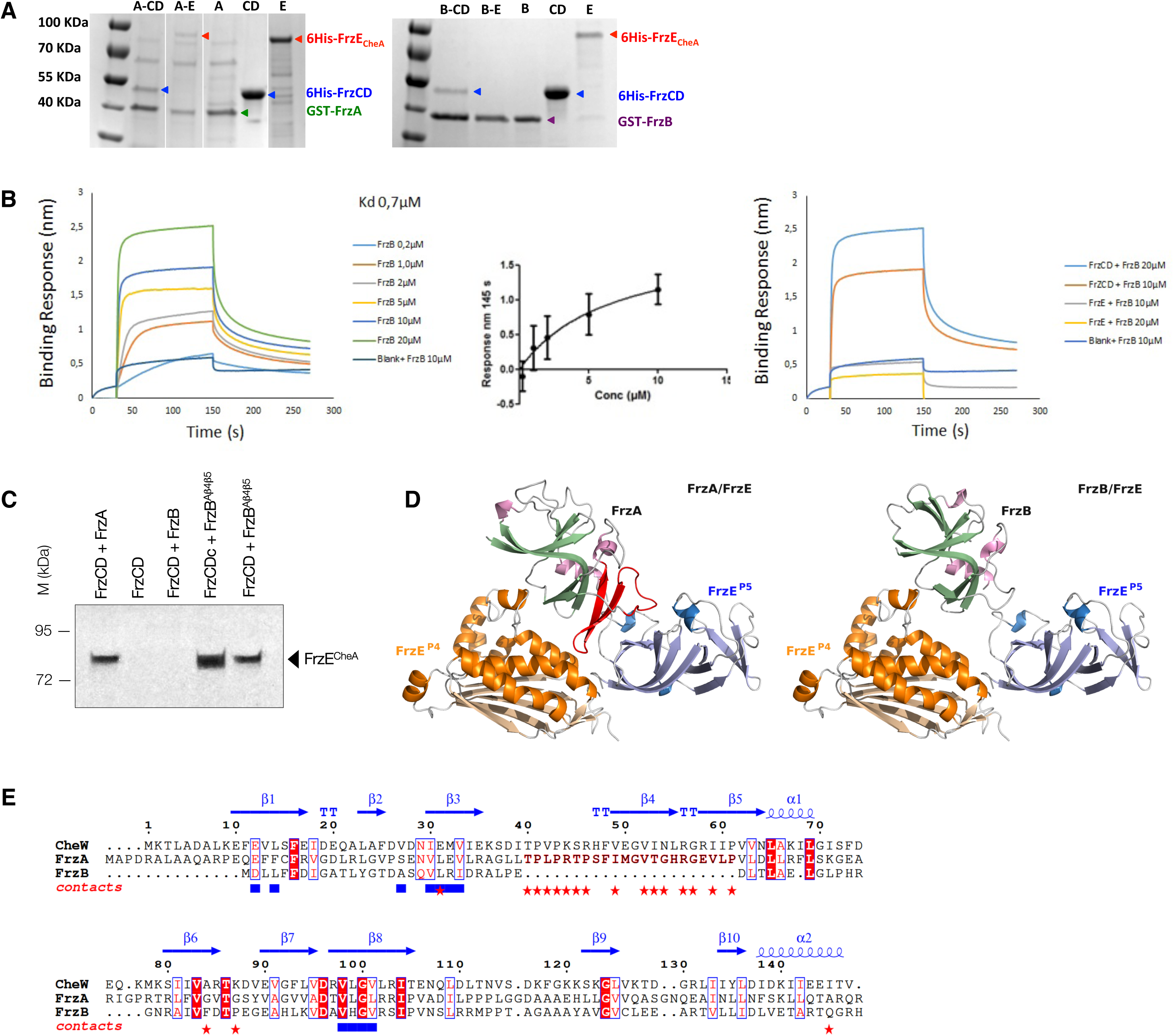
Interaction of FrzB with FrzCD and FrzE. (A) GST affinity chromatography co-purification of the indicated pairs. A, B, CD and E stand for GST-FrzA, GST-FrzB, 6His-FrzCD and 6His-FrzE^CheA^, respectively. In the first gel, lanes are issued from the same gel and were grouped for simplicity. (B) In the left panel, BLi sensograms show the binding of different amounts of GST-FrzB with immobilized 6His-FrzCD. BlitZ ProTM was used to calculate the affinity constant (Kd = 0.7 μM), using a global 1:1 fit. The central panel shows the responses at 145 s (nm, x axis) (average of two biological replicates) of the binding of immobilized 6His-FrzCD with different concentrations of GST-FrzB (μM, y axis). Data were analyzed by GraphPad Prism 5.0, based on steady-state levels of the responses. In the right panel, BLi sensograms show the binding of different amounts of GST-FrzB with immobilized 6His-FrzCD or 6His-FrzE^CheA^ for comparison. A reference subtraction (sensor reference and GST-FrzB at 10 µM) was applied to account for non-specific binding, background, and signal drift to minimize sensor variability. (C) The effect of the FrzB^β4-β5^ chimera on the FrzE phosphorylation. The FrzE kinase domain (FrzE^CheA^) auto-phosphorylation was tested *in vitro* by incubation of FrzE^CheA^ in the presence of FrzA, FrzB or FrzB^β4-β5^, the indicated different forms of FrzCD and ATPγP^33^ as a phosphate donor. (D) 3D models of the FrzE^P4-^ P5:FrzA and FrzE^P4-P5^:FrzB complexes based on the *T. maritima* CheA^P4-P5^:CheW crystal structure (PDB 2ch4). The 20 amino acid region forming the β-strands 4 and 5 of subdomain 2 and corresponding to the CheW region important for the CheW:CheA^P5^ interaction are highlighted in red in the FrzA structure. These residues are missing in FrzB. (E) Protein structural alignments between the *T. maritima* CheW x-ray structure and *M. xanthus* FrzA and FrzB theoretical 3D models. CheW residues in atomic contact with CheA and MCP are shown as red stars and blue squares, respectively. Secondary structures determined from CheW x-ray structure are shown on top of the alignment.

If FrzB does not interact with FrzE, it should not be able to promote the phosphorylation of the FrzE kinase domain (FrzE^CheA^) *in vitro*. In fact, it has been previously shown that the presence of both FrzCD and FrzA is strictly required to achieve the transfer of a phosphoryl group from ATP to the Histidine at position 49 of FrzE^CheA^ *in vitro* (25) (Fig. 3C). As expected, when we replaced FrzA with FrzB in the mixture, no phosphorylation was detected, suggesting that FrzB cannot mediate the FrzE^CheA^ autophosphorylation *in vitro* (Fig. 3C).

Next, we asked what prevented FrzB from interacting with FrzE. To address this question, we decided to generate 3D homology models of the FrzE^P4-P5^:FrzA and FrzE^P4-P5^:FrzB complexes based on the *T. maritima* CheA^P4-P5^:CheW crystal structure (Fig. 3D) (8). The structural models predict that FrzB cannot interact with FrzE due to the lack of a 20 amino acid region forming the β-strands 4 and 5 of subdomain 2. These strands correspond exactly to the CheW region important for the CheW: CheA^P5^ interaction (Fig. 3D and E) (8). On the other hand, and consistent with the two-hybrid system results, FrzB is predicted to contain all the hydrophobic amino acids involved in its interaction with FrzCD (Fig. 3E).

To verify if the lack of the FrzE-interacting region in FrzB was directly associated with the loss of the FrzB-FrzE interaction, we designed a chimeric FrzB protein carrying the β-strands 4 and 5 from FrzA (FrzB^β4-β5^). First, we tested if purified FrzB^β4-β5^ was able to replace FrzA in our FrzE^CheA^ phosphorylation *in vitro* assay. Strikingly, when FrzB^β4-β5^ was the only CheW in the reaction mixture, it promoted the FrzE^CheA^ phosphorylation similarly to FrzA, thus acting as a CheW (Fig. 3C). The FrzE^CheA^ phosphorylation levels were a direct consequence of the FrzCD activation state, because when we used the hyperactive FrzCD^c^ allele in the place of its wild-type version, we observed a higher level of phosphorylated FrzE^CheA^ (Fig. 3C).

Then, to test if FrzB^β4-β5^ could mediate FrzE autophosphorylation *in vivo* as *in vitro*, we transferred the gene coding for FrzB^β4-β5^ into a *ΔfrzA ΔfrzB* strain (Figure S1) at the endogenous locus. FrzB^β4-β5^ was stably expressed (Figure S2), and cells could reverse in the presence of IAA similarly to *ΔfrzB* (Fig. 2). These results suggest that *in vivo*, like *in vitro*, FrzB^β4-β5^ can connect FrzCD to FrzE.

### FrzB might be a stabilizing factor of multiple Frz signaling arrays

It has been previously shown that Frz proteins form nucleoid-associated signaling complexes (26,27). To verify whether FrzB was part of the Frz clusters, we constructed a *mCherry*-*frzB* fusion that replaced *frzB* at the endogenous locus and encoded a functional protein (Figure S2 and S3). mCherry-FrzB formed multiple nucleoid-associated clusters that resembled those formed by FrzCD-GFP and FrzE-mCherry (Fig. 4A) (27).

**Fig. 4.**
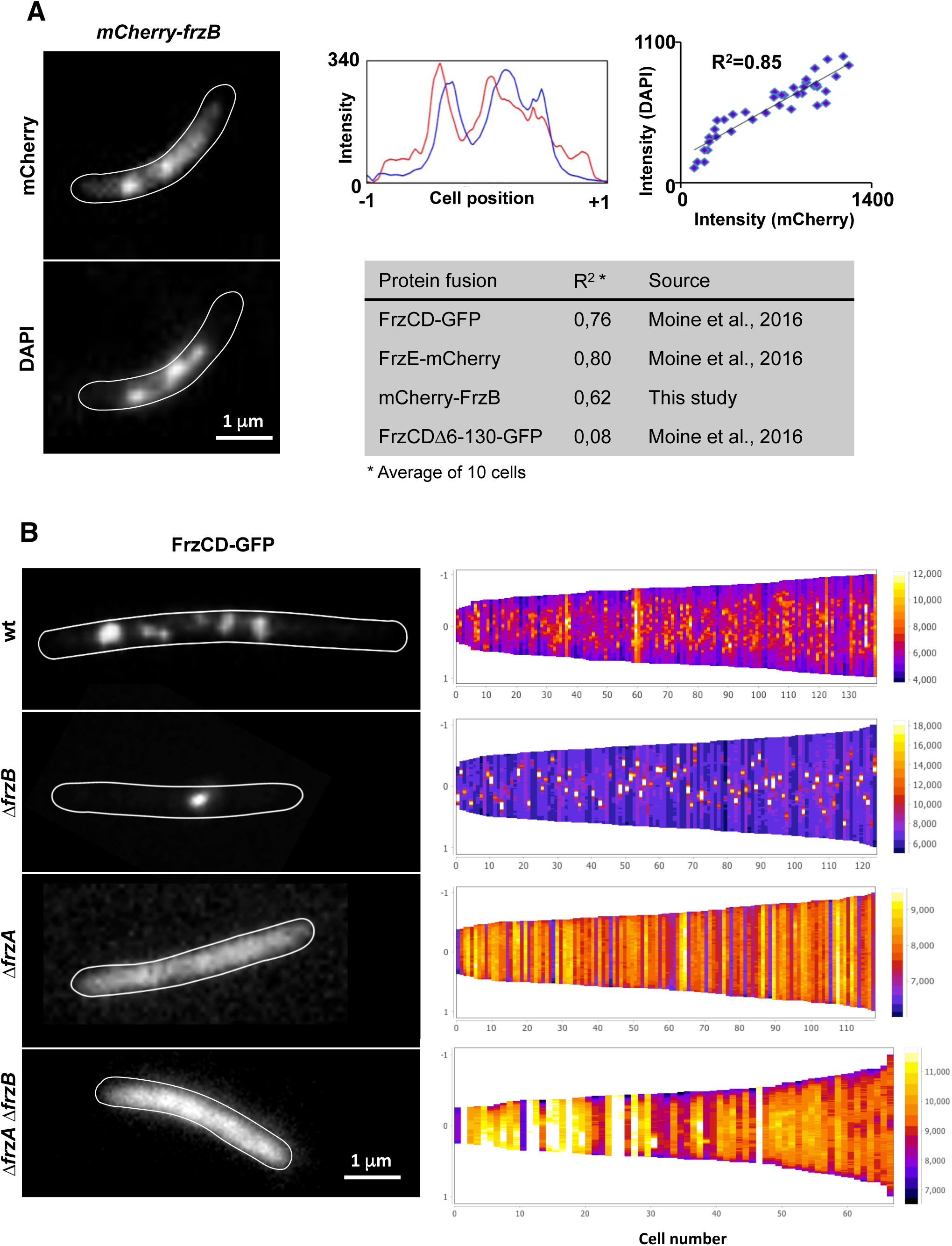
Localization of FrzB-mCherry in *M. xanthus* cells. (A) (Left) Micrographs of *mCherry-frzB* cells stained with the DNA DAPI stain. (Middle) mCherry (red) and DAPI fluorescence (blue) profiles are shown with the fluorescence intensity (arbitrary units) represented on the *y*-axis and the cell length positions with −1 and +1 indicating the poles, on the *x-*axis. The cell boundaries were drawn manually from the phase-contrast images. (Right) Plot of correlation coefficients of DAPI and mCherry localization. R^2^ values > 0.5 indicate significant correlations. The table compares the R^2^ values between the indicated fluorescent fusion and DAPI staining. (B) (Left) Fluorescence micrographs of the indicated *M. xanthus* strains carrying FrzCD-gfp fusions. The cell boundaries were drawn manually from the phase-contrast images. (Right) For each indicated strain, more than 120 cells (*x* axis) from at least two biological replicates are represented as lines and ordered according to their length (pixels) in demographs. The GFP fluorescence intensity along the cell body is represented as colored pixels at the corresponding cell position (from −1 to +1 on the *y* axis). “0” is the cell center. On the right, a scale indicates the fluorescence intensity and the corresponding colors.

Then, we decided to check how the absence of FrzA and FrzB affected the formation of these complexes. For that, we transferred a *frzCD-gfp* gene fusion (27) into *ΔfrzA* and *ΔfrzB* mutants. *M. xanthus* cells of the resulting strains were then imaged by fluorescence microscopy. While FrzCD-GFP was completely dispersed in the cytoplasm in the absence of FrzA, this fusion protein formed a single nucleoid-associated cluster in the absence of FrzB (Fig. 4B). These results suggest that, even if FrzB is dispensable for the assembly of Frz chemosensory arrays, it is important for the formation of multiple distributed clusters on the nucleoid. These clusters collapse into a single cluster in the absence of FrzB. Of 230 cells examined, 82% have a single cluster, whereas 18% have no cluster. It is possible that, in the absence of FrzB and during cell division, only one daughter cell inherits the single Frz array and the other one will synthesize a new one. This would explain the lack of clusters in a small population of cells. In the wild type, the multiple Frz clusters would be expected to be distributed to both daughter cells (27). In the absence of both FrzA and FrzB, FrzCD-GFP was diffused in the cytoplasm, as in *ΔfrzA* (Fig. 4B). The aberrant localization patterns were not due to a change in the FrzCD-GFP protein levels nor in a change of the nucleoid structure (Figure S5 and Figure S6). Interestingly, and by yet unknown mechanisms, FrzA not only is essential for Frz cluster formation, but it also allows the association of FrzCD to the nucleoid (Fig. 4).

## DISCUSSION

Chemosensory systems are modified two-component systems in which the addition of accessory proteins confers unique properties like signal amplification and sensitivity. In this work, we found that the CheW-like protein FrzB lacks a 20 amino acid sequence corresponding to two β-strands that are important for CheA^P5^:CheW interactions (8). Thus, FrzB cannot interact with FrzE and mediate its autophosphorylation. The loss of this protein region might have been selected to confer to this small protein the unique function of allowing the formation and distribution of multiple nucleoid Frz chemosensory arrays. How would FrzB accomplish these tasks? It has been shown that Che arrays are constituted by hexagons of MCP trimers of dimers networked by CheA^P5^-CheW rings (11). By homology, we believe that Frz proteins can have such hexagonal organization using the nucleoid as scaffold. However, if FrzB cannot interact with FrzE^P5^, it cannot participate to the formation of CheA^P5^-CheW rings either. In addition to CheA^P5^-CheW rings, the formation of Che networks implies the formation of six-member CheW rings that only interact with the MCPs (Fig. 1) (17,28,29). The 6-CheW rings have been visualized by cryoEM, and their suggested function is to stabilize the Che arrays by favoring allosteric interactions, like CheW-CheA rings, but without a direct effect on signaling. We can imagine that 6-FrzB rings (in addition to FrzE^P5^-FrzA rings and 6-FrzA rings) might further stabilize small Frz clusters, which would otherwise collapse into one single cluster (Fig. 5). Alternatively FrzB allows the formation of small Frz clusters by breaking the networking of MCP trimers of dimers with Che^P5^-CheW rings (Fig. 5).

**Fig. 5.**
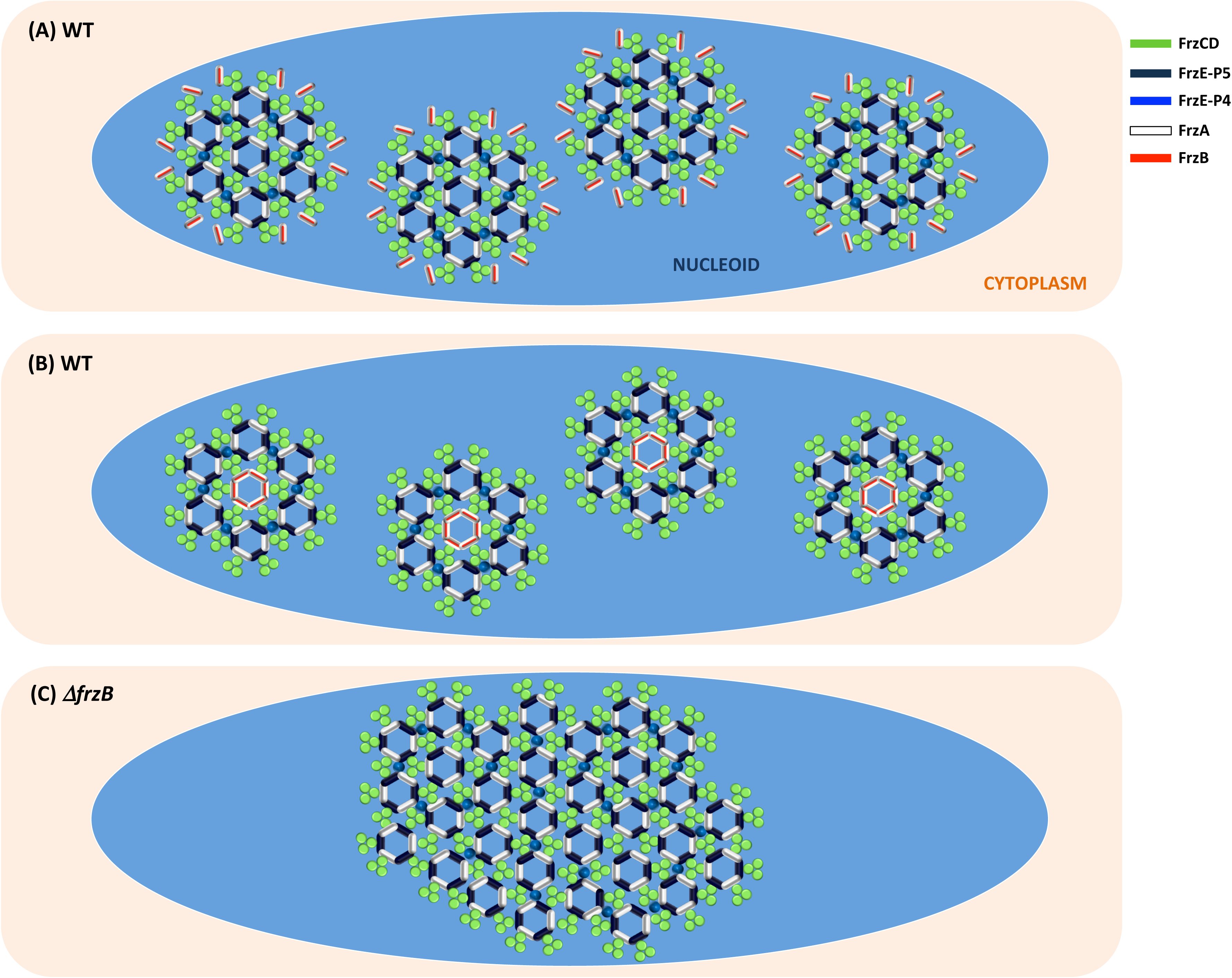
Proposed model on how FrzB might confer plasticity to Frz clusters. (A) FrzB participates to the formation of 6-CheW rings to stabilize small Frz clusters, which would otherwise collapse in one single cluster (C). (B) Alternatively FrzB allows the formation of small Frz clusters by breaking the networking of MCP trimers of dimers with Che^P5^-CheW rings.

Interestingly, the FrzCD receptor does not contain an obvious ligand-binding domain. Instead, it contains an N-terminal region that has been recently shown to be involved in DNA binding (27). Thus, FrzB could also function to deliver signals to the Frz pathway. FrzB might bind directly to some cytoplasmic molecules and transduce signals to FrzCD, thus functioning as a sensor of the cellular metabolic state. However, FrzB could also connect FrzCD to another MCP or sensor kinase. *M. xanthus* encodes a total of 21 MCPs (30). Among them, only Mcp7 is cytoplasmic like FrzCD, but its localization in a single cluster randomly localized within cells, strongly excludes an association with the Frz pathway (30). We could imagine that FrzCD, which forms clusters on the nucleoid (27), could associate with a transmembrane MCP or sensor kinase at regions where the nucleoid surface is particularly close to the inner membrane (31). In fact, previous work shows the association of FrzCD with membrane fractions (32) as well as interactions with envelope-associated A-motility proteins (33,34). The hypothesis that FrzB is a Frz input does not rule out that FrzCD itself can somehow bind signals. Indeed, FrzCD responds to IAA independently of FrzB (Fig. 2).

In recent years, the discovery of different accessory Che proteins allowed the identification of new functions and degrees of complexity in the regulation of the cellular responses operated by chemosensory systems. By characterizing a new accessory protein and its function, this work opens new insights on the knowledge of the regulatory potentials of bacterial chemosensory systems.

## MATERIALS AND METHODS

### Bacterial strains, plasmids and growth

*S*trains and plasmids are listed in Table S1 and S2, respectively. *M. xanthus* strains were grown at 32°C in CYE rich media as previously described.

Plasmids were introduced into *M. xanthus* cells by electroporation. Deletions and GFP fusions were inserted in frame, using the pBJ113 or pBJ114 vectors, to avoid polar effects on the downstream gene expression (22,30).

Strains EM736 and EM743 were obtained by deleting *frzA* from codon 15 to 145 and *frzB* from codon 11 to 92, respectively. In both strains we also deleted *frzCD* from codon 6 to 182. Strain EM742 was obtained by deleting *frzA* and *frzB* as above.

To construct plasmid pEM399 to generate EM515, a DNA fragment was generated by PCR overlapping a DNA region amplified with primers CCCAAGCTTCTTCAGCTCGAGGCTGGCAA and TGCCCCTTGCTCACCATCTCCTCCTCCTTCTCATGGAA and *mCherry* amplified with primers CATGAGAAGGAGGAGGAGATGGTGAGCAAGGGCGAGGA and CGGGATCCTCTAGACTTGCACAGCTCGTCCATGC from pEM147 (33). This fusion product was digested with HindIII and BamHI. *frzB* was amplified with primers CGGGATCCGGAGCAGGAGACCTCCTCTTCTTCGACAT and GGAATTCGCGGACGAGGACAGCCTCAA and digested with BamHI and EcoRI. The two fragments were then cloned into pBJ113 previously digested with HindIII and BamHI in a three-partner ligation reaction.

To obtain strains EM440 and EM512, plasmid pDPA20 was transferred into strains DZ4479 and DZ4478 respectively.

For the construction of *frzB*^*β4-β5*^ we replaced codon 27 to 36 of FrzB with codon 41 to 75 of FrzA. Plasmids pEM463 and was synthesized by Biomatik.

*Escherichia coli* cells were grown under standard laboratory conditions in LB broth supplemented with antibiotics if necessary.

### Phenotypic analysis

Developmental assays were performed by spotting 10 μl of cells at 5 OD_600_ directly onto CF 1.5 % agar plates. Vegetative swarming phenotypes were analyzed by spotting the same amount of cells onto CYE plates containing a concentration of 0.5 % agar. The IAA response of cell colonies was analyzed by spotting cells (10 μl of cells at 5 OD_600_) on CYE plates supplied with 0.15 % of IAA and containing a concentration of 1.5% agar. After incubation at 32°C the swarming colonies and the fruiting bodies were photographed at an Olympus SZ61 stereoscope with a Nikon DXM1200 Digital Camera.

### Reversal frequencies

5 μl of cells from 4 × 10^8^ cfu ml^−1^ vegetative CYE cultures were spotted on a thin fresh TPM agar supplied or not with 0.15 % IAA. Time-lapse movies were shot for 1 hour with frames captured every 30 seconds. Movies were obtained and reversal frequencies calculated with Image J (Rasband, W.S., ImageJ, U. S. National Institutes of Health, Bethesda, Maryland, USA, http://imagej.nih.gov/ij/, 1997-2012) and FIJI as previously described (20). For non-reversing strains, the number of reversals for each cell was plotted against time using the R software (https://www.r-project.org/). For strains that frequently reversed, the number of periods between two reversals were plotted against time using the R software. Reversal frequencies were measured from cells issued from two to six biological replicates. Statistical significance was obtained with a Wilcox Test from R.

### Protein purification and co-purification

BL21(DE3) [F– ompT hsdSB(rB– mB–) gal dcm (DE3)] cells were grown in LB broth supplemented 100 µg/ml ampicillin to mid-exponential phase at 37°C. Overexpression was induced by adding 0,2 mM IPTG. Cells were then grown at 16°C over night. Cells were washed and resuspended in lysis buffer (50 mM TrisHCl, pH 8; 300 mM NaCl; 100 µg/ml PMSF; 30 U/mL Benzonase) and lysed at the French press. The cell lysates were centrifuged at 4°C for 20 min at 18000× rpm.

For BLi experiments, soluble 6His-tagged proteins were purified using a NiNTA resine (GE Healthcare) and imidazole was removed by dialysis into a buffer containing 50 mM TrisHCl, pH 8 and 300 mM NaCl. GST-proteins were purified by GST-affinity chromatography using the same lysing buffer supplied of 10 mM reduced glutathione to elute.

For co-purification experiments, *E. coli* cells expressing each of GST-FrzA, GST-FrzB, 6His-FrzCD or 6His-FrzECheA were mixed at one to one ratio to obtain the following combinations: GST-FrzA: 6His-FrzCD, GST-FrzA: 6His-FrzECheA, GST-FrzB: 6His-FrzCD, GST-FrzB: 6His-FrzECheA. Mixed cell cultures were resuspended in lysis buffer (50 mM TrisHCl, pH 8; 300 mM NaCl; 100 µg/ml PMSF; 30 U/mL Benzonase) and lysed at the French press. The cell lysates were centrifuged at 4°C for 20 min at 18000× rpm.

Proteins were, then, co-purified by GST-affinity chromatography as described above.

### Biolayer Interpherometry

Protein-protein interaction experiments were conducted at 25°C with the BLItz instrument from ForteBio (Menlo Park, CA, USA). The BLI consists in a real time optical biosensing technique exploits the interference pattern of white light reflected from two surfaces to measure biomolecular interactions (24). Purified 6His-FrzCD or 6His-FrzE^CheA^ protein ligands were immobilized onto two different Ni-NTA biosensors (ForteBio) in duplicate at 4 µM concentrations. GST-FrzB was used as the analyte throughout the study at concentrations ranging from 0.2 to 20 µM. The assay was conducted in buffer containing Tris HCl 50 mM pH8 and NaCl 300 mM. The binding reactions were performed with an initial baseline during 30 seconds, an association step at 120 seconds and a dissociation step of 120 seconds with lateral shaking at 2200rpm. A double reference subtraction (sensor reference and GST-FrzB at 10 µM) was applied to account for non-specific binding, background, and signal drift to minimize sensor variability.

### Sequence alignment and homology modeling

Predictions of secondary structures and protein sequence alignments were obtained with Jpred (35) and Clustal Omega (36), respectively. Homology models of FrzA (Uniprot P43498) or FrzB (Uniprot P43499) in the presence of FrzE (Uniprot P18769) were built with MODELLER 9v12 (37). Crystal structure of the complex between bacterial chemotaxis histidine kinase CheA^P4-P5^ and receptor-adaptor protein CheW (PDB codes 2CH4) was used as template (38). The modeling was performed in a two-step manner: First, each individual domain was modeled separately using the 3D structure of its *T. maritima* counterpart as a template. CheW (2CH4 chain W) was used to build models of FrzA and FrzB. The N-terminal domain of CheA (2CH4 chain A resid 355-540) and the C-terminal domain of CheA (2CH4 chain A resid 541-671) were used to build N- and C-terminal domains of FrzE. In a second step, each model was superimposed to its counterpart in the x-ray structure of the CheW-CheA complex (chains W and A). The Frz proteins exhibit high enough identities to their Che counterparts to build reasonable 3D models. FrzB vs CheW (26.1% identity, 45% similarity), FrzA vs CheW (26.6% identity, 46.7% similarity), FrzE^p4^ vs. CheA^p4^ (46.5% identity, 71.7% similarity), FrzE^p5^ vs. CheA^p5^ (26.1% identity, 53.8% similarity). Quality of the models was assessed with Prosess server (http://www.prosess.ca). Structural alignments were performed with Protein structure comparison service PDBeFold at European Bioinformatics Institute (http://www.ebi.ac.uk/msd-srv/ssm) (39). Fig. 3C showing the lowest energy model was generated with pymol (http://pymol.org). Fig. 3D was generated with ESPript web server (40).

### Fluorescence microscopy and image analysis

For fluorescence microscopy analyses, 5 μl of cells from 4 x 10^8^ cfu ml^−1^ vegetative CYE cultures were spotted on a thin fresh TPM agar pad at the top a slide (41). A cover slip was added immediately on the top of the pad, and the obtained slide was analyzed by microscopy using a Nikon Eclipse TE2000 E PFS inverted epifluorescence microscope (100 x oil objective NA 1.3 Phase Contrast) (42).

To study the co-localization with the DNA, the TPM agar pads were supplied with 1µg ml^−1^ DAPI stain as shown by Moine et al. (27). FrzB clusters, nucleoid areas and cell areas were automatically detected and verified manually with the “MicrobeJ” Fiji/ImageJ plugin created by A. Ducret, Brun Lab (http://www.indiana.edu/~microbej/index.html). All data plots and statistical tests were obtained with the R software (https://www.r-project.org/).

### *In vitro* autophosphorylation assay

*In vitro* phosphorylation assays were performed with *E. coli* purified recombinant proteins. 4 μg of FrzE^CheA^ was incubated with 1 μg of FrzA, FrzB or FrzB^β4-β5^ and FrzCD or FrzCD^c^ in 25 μl of buffer P (50 mM Tris-HCl, pH 7.5; 1 mM DTT; 5 mM MgCl_2_; 50 mM KCl; 5 mM EDTA; 50 μM ATP, 10% (v/v) glycerol) supplemented with 200 μCi ml^−1^ (65 nM) of [γ-33P]ATP (PerkinElmer, 3000 Ci mmol^−1^) for 10 minutes at room temperature in order to obtain the optimal FrzE^CheA^ autophosphorylation activity. Each reaction mixture was stopped by addition of 5 × Laemmli and quickly loaded onto SDS-PAGE gel. After electrophoresis, proteins were revealed using Coomassie Brilliant Blue before gel drying. Radioactive proteins were visualized by autoradiography using direct exposure to film (Carestream).

## Acknowledgements

We thank Romain Mercier, Dorothée Murat and Laetitia My for their support and helpful discussions. We also thank Anne-Valerie Le Gall for her technical support. We thank the anonymous reviewers, whose comments helped to greatly improve the manuscript.

## SUPPLEMENTARY FIGURE AND TABLE LEGEND

**Figure S1.**
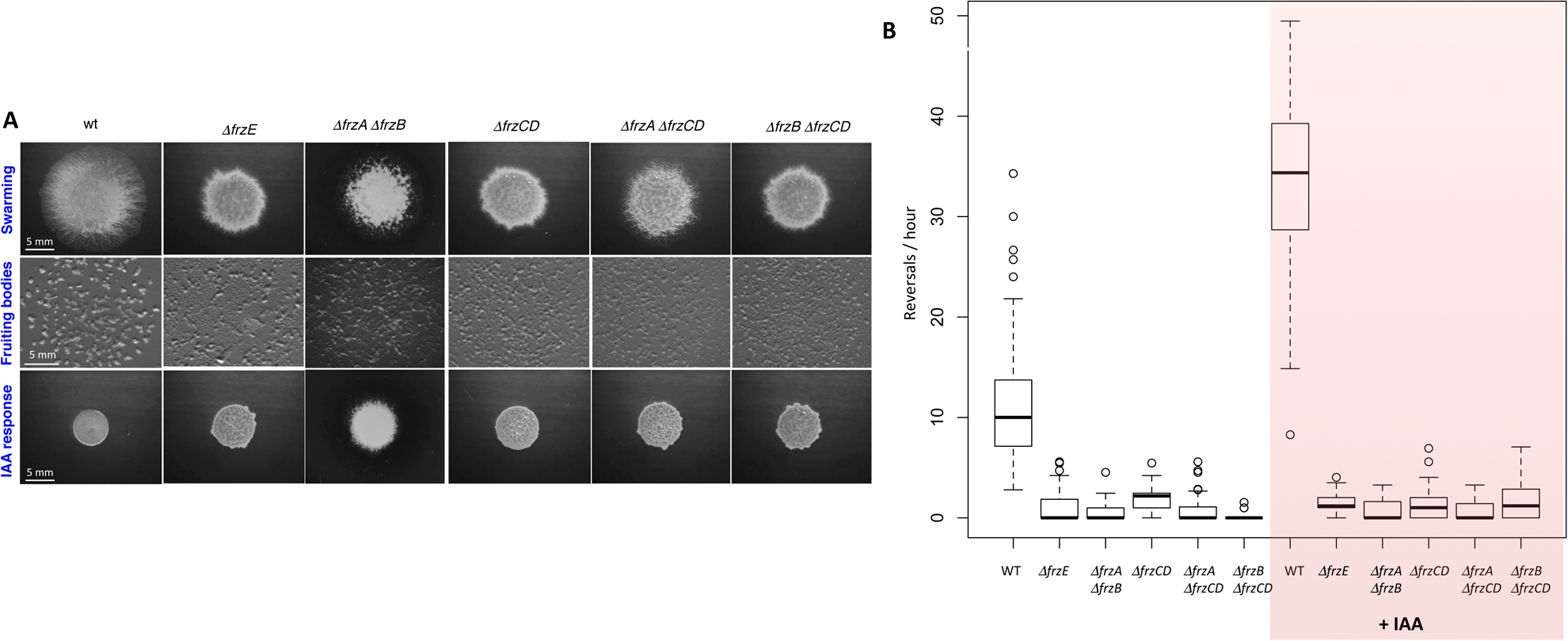
(A) Motility and fruiting body formation phenotypes were photographed at 48h and 72h, respectively. (B) Box plots of reversal frequencies of single cells moving on agar pads supplemented or not with 0.15% IAA. The lower and upper boundaries of the boxes correspond to 25% and 75% percentiles, respectively. The median is shown as a line at the center of each box and whiskers represents the 10% and 90% percentiles. For the reversal frequency measurements, approximately 120 cells, issued from two biological replicates, were used. For WT approximately 500 cells, issued from six biological replicates, were used.

**Figure S2.**
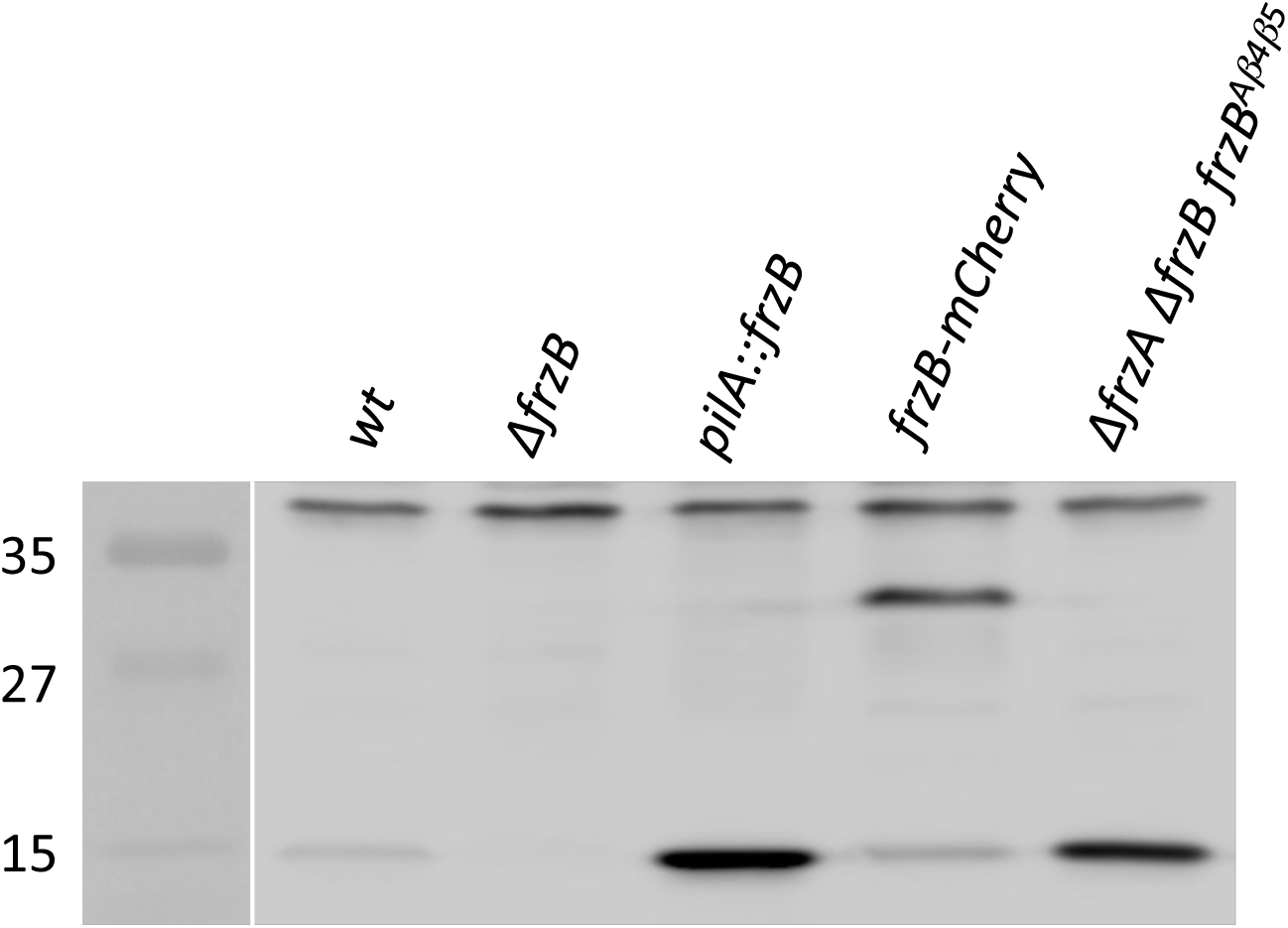
Western blot with anti-FrzB antibodies on the cell extracts of the indicated *M. xanthus* strains. The *pilA::frzB* strain, used as positive control, expresses *frzB* under the control of the *pilA* promoter for high expression. The white line is used to indicate that two lanes from the same gel were separated by other lanes in the original gel.

**Figure S3.**
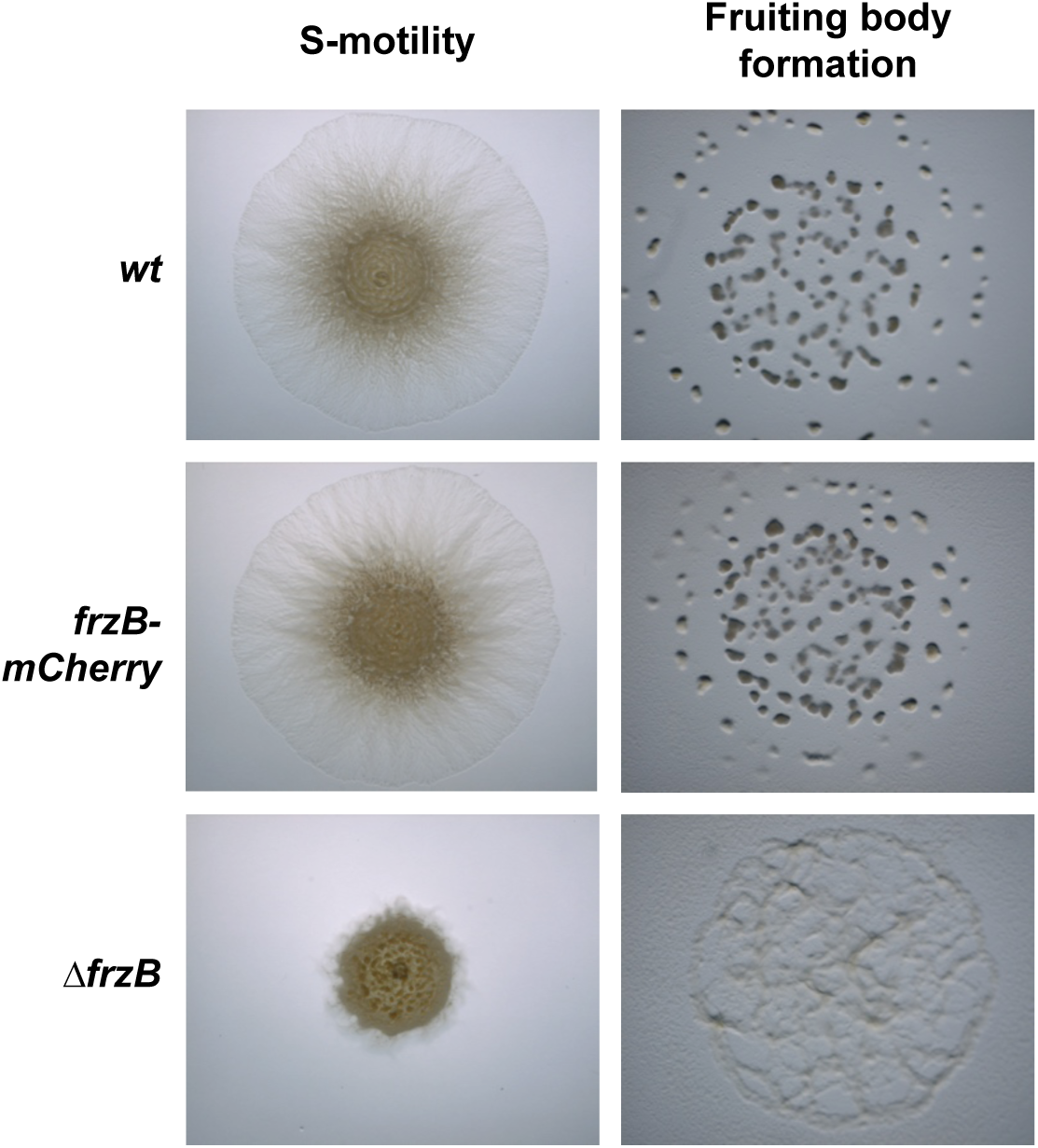
Motility and fruiting body formation phenotypes of the indicated strains were photographed at 48h and 72h, respectively.

**Figure S4.**
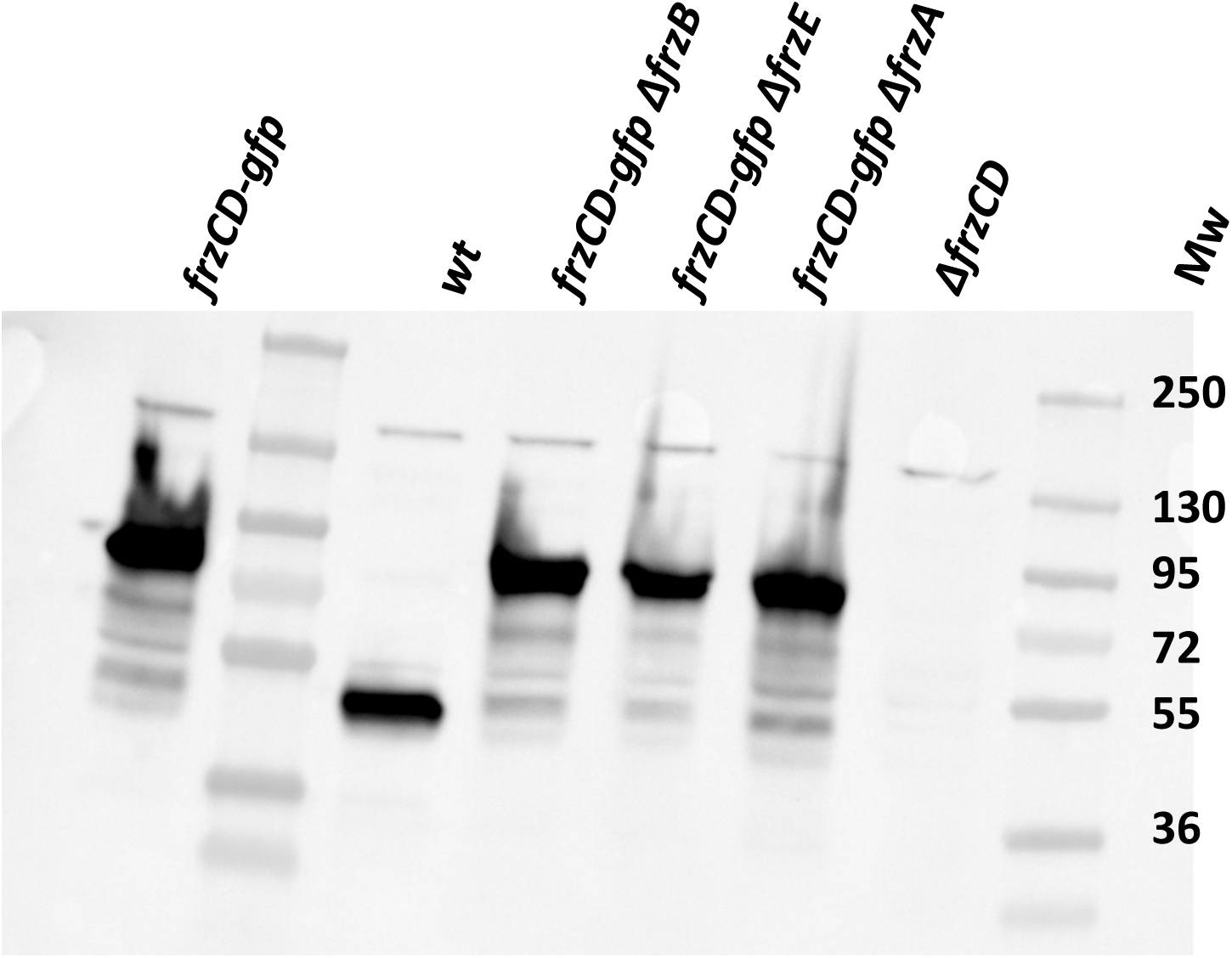
Western blot with anti-FrzCD antibodies on the cell extracts of the indicated *M. xanthus* strains.

**Figure S5.**
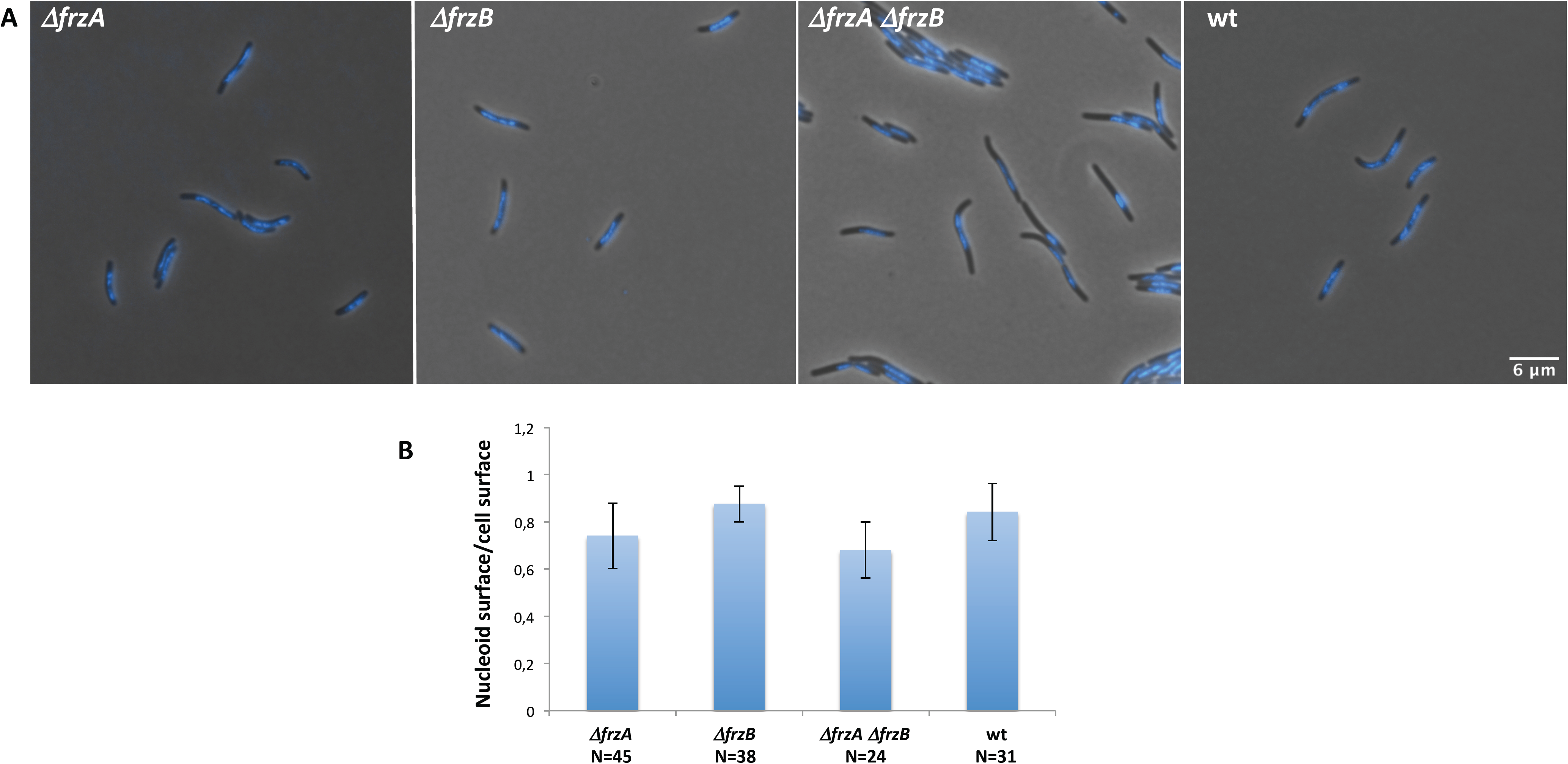
(A) Cells of the indicated strains were incubated 30 minutes with DAPI and then imaged at the fluorescence microscope. The nucleoid and cell surfaces were measured automatically with Microbe J (44). (B) The ratio between the nucleoid and cell surface was then calculated for each cell and averages values plotted. The numbers of analyzed cells are indicated per each strain. Cells were issued from two independent biological replicates.

**Table S1.** List of strains used in this work.

**Table S2.** List of plasmids used in this work.

**Table S1.** Strain list.

| Strains | Genotype | Reference or source |
| --- | --- | --- |
| DZ2 | *wt* | Zusman et al., 1982 |
| DZ4478 | Δ*frzA* | Bustamante et al ; 2004 |
| DZ4479 | Δ*frzB* | Bustamante et al ; 2004 |
| DZ4487 | *frzCD*Δ6-182 | Bustamante et al ; 2004 |
| DZ4480 | Δ*frzCD* | Bustamante et al ; 2004 |
| EM538 | Δ*frzA frzCD*Δ6-182 | This study |
| EM539 | Δ*frzB frzCD*Δ6-182 | This study |
| DZ4620 | *frzCD-gfp* | Mauriello et al., 2009 |
| EM440 | Δ*frzB frzCD-gfp* | This study |
| EM512 | Δ*frzA frzCD-gfp* | This study |
| EM515 | *frzB-mCherry* | This study |
| EM605 | Δ*frzA* Δ*frzB frzB*<sup>β4-β5</sup> | This study |
| EM736 | Δ*frzA* Δ*frzCD* | This study |
| EM742 | Δ*frzA* Δ*frzB* | This study |
| EM743 | Δ*frzB* Δ*frzCD* | This study |
| EM699 | Δ*frzB frzB*<sup>β4-β5</sup> | This study |
| EM700 | Δ*frzA frzB*<sup>β4-β5</sup> | This study |
| EM773 | Δ*frzA* Δ*frzB frzCD-gfp* | This study |

**Table S2.** Plasmid list.

| Plasmid | Expression plasmid | Source |
| --- | --- | --- |
| pEM411 | pBJ113 with construct for $\Delta frzA frzCD\Delta 6-182$ | This study |
| pEM412 | pBJ113 with construct for $\Delta frzB frzCD\Delta 6-182$ | This study |
| pEM463 | pBJ113 with construct $\Delta frzA \Delta frzB frzB^{\beta 4-\beta 5}$ (Biomatik) | This study |
| pEM524 | pBJ114 with construct $\Delta frzA \Delta frzB$ | This study |
| pEM525 | pBJ114 with construct $\Delta frzA$ in DZ4480 | This study |
| pEM526 | pBJ114 with construct $\Delta frzB \Delta frzCD$ | This study |
| pEM536 | pBJ114 with construct $\Delta frzA frzB^{\beta 4-\beta 5}$ | |
| pEM537 | pBJ114 with construct $\Delta frzB frzB^{\beta 4-\beta 5}$ | |
| pDPA20 | pKY480 with construct for $frzCD-gfp$ | Mauriello et al., 2009 |
| pEM399 | pBJ113 with construct for $frzB-mCherry$ | This study |
| pETPhos $frzCD$ | pETPhos with $frzCD$ tagged with 6-his inducible with IPTG | Guzzo et al., 2015 |
| pETPhos $frzE^{CheA}$ | pETPhos with $frzE^{kinase}$ tagged with 6-his inducible with IPTG | Guzzo et al., 2015 |
| pETPhos $frzCD^c$ | pETPhos with $frzCD^c$ tagged with 6-his inducible with IPTG | Guzzo et al., 2015 |
| pGEX(M) $frzA$ | pGEX (M) with $frzA$ tagged with GST | Guzzo et al., 2015 |
| pEM464 | pETPhos with $frzB^{\beta 4-\beta 5}$ tagged with 6-his inducible with IPTG (Biomatik) | This study |

